# Exo-Enzymatic Cell-Surface Glycan Labeling for Capturing Glycan-Protein Interactions through Photo-Crosslinking

**DOI:** 10.1101/2022.01.20.477088

**Authors:** Jonathan L. Babulic, Chantelle J. Capicciotti

**Affiliations:** Department of Biomedical and Molecular Sciences, Queen’s University, Kingston, K7L 3N6, Canada; Department of Chemistry, and Department of Surgery, Queen’s University, Kingston, K7L 3N6, Canada

**Keywords:** Exo-enzymatic labeling, Glyco-engineering, sialyltransferases, photo-crosslinking sugars, sugar-nucleotide derivatives, chemo-enzymatic synthesis

## Abstract

Tools to interrogate glycoconjugate-protein interactions in the context living cells are highly attractive for the identification of critically important functional binding partners of glycan-binding proteins. These interactions are challenging to interrogate due to low affinity and rapid dissociation rates of glycan-protein binding events. The use of photo-crosslinkers to capture glycan-protein interaction complexes has shown great promise for identifying binding partners involved in these interactions. Current methodologies use metabolic oligosaccharide engineering (MOE) to incorporate photo-crosslinking sugars. However, these MOE strategies are not amenable to all cell types and can result in low incorporation and cell-surface display of the photo-crosslinking probe, limiting their utility for studying many types of interactions. We describe here an exo-enzymatic strategy for selectively introducing photo-crosslinking probes into cell-surface glycoconjugates using the recombinant human sialyltransferase ST6GAL1 and a diazirine-linked CMP-Neu5Ac derivative. Probe introduction is highly efficient, amenable to different cell types and resulted in improved crosslinking when compared to MOE. This exo-enzymatic labeling approach can selectively introduce the photo-crosslinking sugar on to specific glycan epitopes and subclasses by harnessing the specificity of the sialyltransferase employed, underscoring its potential as a tool to interrogate and identify glycoconjugate ligands for diverse glycan-binding proteins.

## Introduction

Cell-surface glycans and glycoconjugates play essential roles in the coordination of many physiological processes.^1^ Glycans are central to cell-cell and cell-matrix signaling interactions, often through highly specific recognition events mediated by glycan-binding proteins. For example, glycan recognition by galectins or Siglecs are involved in immune recognition, selectin-glycan interactions mediate immune cell adhesion to endothelium to permit entry to sites of inflammation, and glycans are widely involved in host-pathogen interactions.^1-4^ Despite their biological importance, interrogating and characterizing these native binding events remains challenging.^5-7^ Glycan-protein interactions are typically transient and low-affinity binding events and have association constants in the millimolar-micromolar range.^8^ In a physiological environment, multivalent glycan/glycoprotein-ligand and protein-receptor presentation strengthen these interactions,^8-11^ however, identifying protein binding partners of glycans necessitates harsh cell lysis which can dissociate these interactions.

To address these challenges, photo-crosslinking probes have been introduced into cellular glycans to capture glycan-protein complexes involved in native cellular binding events. Linkers bearing photoactivatable crosslinking functional groups have been conjugated to monosaccharides for incorporation into cellular glycoconjugates through metabolic oligosaccharide engineering (MOE).^12-20^ Activation of the photo-crosslinking moiety by UV irradiation produces a reactive species that induces formation of a proximity-mediated covalent bond between the glycan and interacting binding partner, allowing for these interactions to be captured and identified. Photo-crosslinking sugars include aryl azide-linked monosaccharides^12-14^ and smaller, more versatile and efficient diazirine-linked sugar analogues developed by the Kohler group.^15, 18, 20^ Diazirine-conjugated sialic acids have been incorporated into cellular glycoproteins and glycolipids through MOE using mannosamine or sialic acid derivatives as unnatural biosynthetic precursors.^15-18, 20, 21^

While MOE is a powerful approach for incorporation of photo-crosslinking and other chemical reporter probes,^12^ it is limited by the tolerance of the appended probe by cellular metabolic machinery. Incorporation efficiency of probe-modified monosaccharides is highly variable and many cell types, including primary cells, are unable to incorporate these probes, limiting the interactions that can be studied.^16, 21-23^ Sugar probes used for MOE can also exhibit toxicity or inhibit the growth of cells,^21, 24^ and there is evidence that commonly used membrane-permeable monosaccharides may yield false positive signals through artificial non-enzymatic cysteine *S-*glycosylation.^25^ Finally, MOE approaches are not amenable to selective installation of photo-crosslinking or other chemical reporter probes as there is no selectivity for glycan subclass- or epitope-specific display, resulting in global incorporation by α-2,6 and α-2,3 sialyltransferases.^21^

Selective exogenous (exo)-enzymatic labeling (SEEL) is a promising alternative strategy to overcome limitations of MOE for remodeling cell-surface glycans. In this approach, modified sugar nucleotides are used with recombinant glycosyltransferase enzymes to install these sugar analogues onto cell-surface glycoconjugates. The SEEL methodology has been applied to install CMP-Neu5Ac derivatives bearing smaller chemical reporters (e.g. azides, alkynes, biotin) and larger biomolecules, including glycans, on to cell-surfaces, predominantly using the sialyltransferase ST6GAL1.^26-32^ There are several advantages to exo-enzymatic labeling, such as high efficiency of labeling, decreased labeling reaction time, and more control over the density display of the chemical probe.^26-28^ Furthermore, this method can be used to selectively install a modified sialic acid with a high degree of linkage and glycan subclass specificity, depending on the sialyltransferase used. For example, the native specificities of human sialyltransferases have been exploited to selectively install azide- or biotin-modified CMP-Neu5Ac on *N*-linked glycans with an α-2,6-linkage using ST6GAL1, and on *O*-linked glycans with an α-2,3-linkage using ST3GAL1.^14, 27, 28, 30, 32-34^ Selective labeling of glycoproteins bearing the sialylated-Thomsen-Friedenreich antigen (sialyl-T) has also been demonstrated using ST6GALNAC4 and a biotin-linked CMP-Neu5Ac.^31^

Herein, we report a broadly applicable methodology for cell-surface display of a diazirine-linked sugar derivative on cellular glycans based on exo-enzymatic transfer of an unnatural CMP-Neu5Ac derivative (Figure 1). In this approach, native cellular glycans are remodeled by first cleaving native sialic acid with a sialidase, followed by selective capping of *N*-glycans with a diazirine (DAz)-conjugated CMP-Neu5Ac derivative (CMP-Neu5DAz, **1**) using the recombinant sialyltransferase ST6GAL1.^26-28, 30, 33^ As this method does not rely on native cellular machinery, this approach is amenable to any cell type and can be used to study interactions inaccessible by MOE probe installation. Furthermore, enzymatic glycan labeling can occur in 2 hours and is highly efficient, circumventing the long incubation times needed for metabolic labeling studies. As a proof-of-principle, cells expressing CD22 (Siglec-2) were labeled with CMP-Neu5DAz (**1**) and CD22 *cis*-interaction (same-cell) complexes were crosslinked *in situ* by UV irradiation. We demonstrate that enzymatic labeling using CMP-Neu5DAz and ST6GAL1 is applicable to multiple cell lines and that density of the probe can be controlled. Furthermore, we show that our approach has enhanced crosslinking efficiency when compared to MOE, demonstrating its potential as a powerful tool for interrogating glycan/protein interactions.

**Figure 1.**
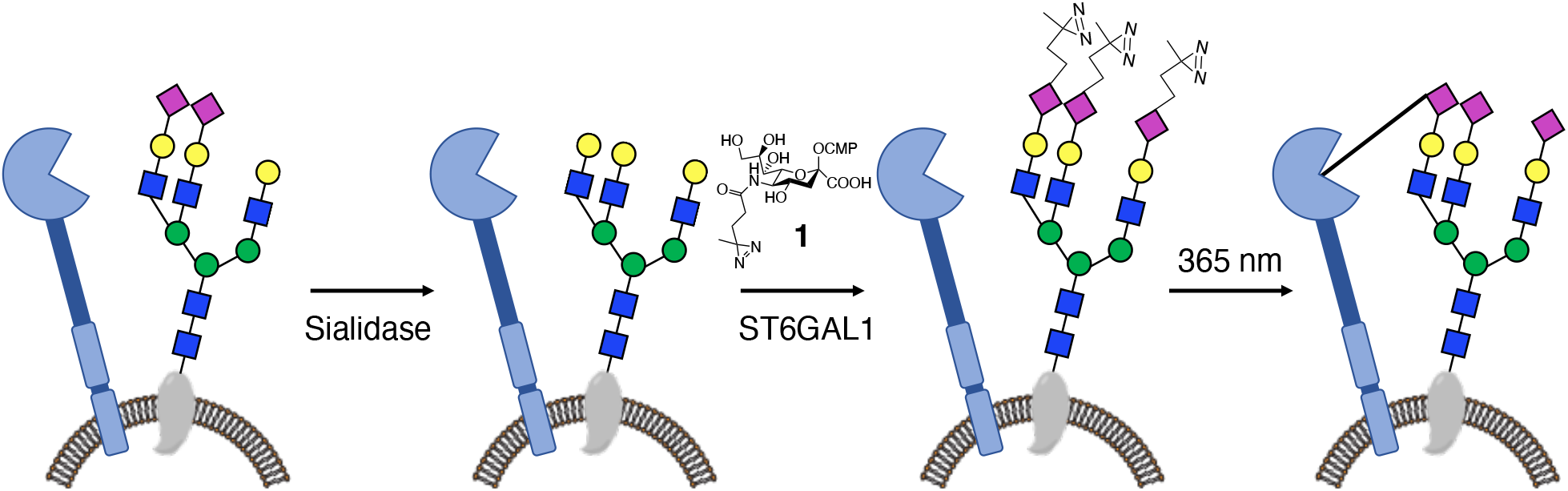
Photo-crosslinking of *cis*-glycan-protein interactions through enzymatic cell-surface glycan labeling using an exogenously administered CMP-Neu5DAz (1) derivative and recombinant ST6GAL1.

## Results and Discussion

### Chemo-Enzymatic Synthesis of CMP-Neu5DAz

The sialyltransferase ST6GAL1 is a promiscuous enzyme that can tolerate a wide range of modifications at the C-5 position of CMP-sialic acid (CMP-Neu5Ac) and this enzyme has been used in cell-surface glycan engineering applications.^26-28^ Preliminary work from the Paulson group also reported enzymatic labeling of cellular glycans using a C-9 aryl azide-linked photo-crosslinking CMP-Neu5Ac analogue.^14^ Access to C-9 CMP-Neu5Ac derivatives is challenging as modification of the C-6 position of the mannosamine precursor is often poorly tolerated by enzymes used for their chemo-enzymatic synthesis.^33, 35^ Furthermore, derivatization at the C-9 position can greatly impact the affinity and recognition of glycan-binding proteins for sialic acid and crosslinked complexes formed using aryl azides may not accurately represent native binding interactions.^15, 36, 37^ A diazirine modified levulinic acid is an attractive photo-crosslinker for the studying of glycan-protein interactions because of its minimal size and relatively unbiased reactivity.^15, 38^ We thus sought to employ a CMP-Neu5Ac derivative bearing a diazirine photo-crosslinking functionality at the C-5 position to engineer cellular glycans for interrogation of glycan-protein interactions.

Simple CMP-Neu5Ac derivatives can be prepared chemo-enzymatically from analogues of their native biosynthetic precursor, *N*-acetylmannosamine (ManNAc).^35^ While various derivatives have been prepared used this strategy, diazirine-modified CMP-Neu5Ac analogues have not been reported. To access a photo-crosslinking CMP-Neu5Ac for exogenous cell-surface labeling, we first prepared a diazirine (DAz) modified levulinic acid linker,^15, 39^ which was conjugated to mannosamine using EDC•HCl, HOBt and Et_3_N to afford a DAz-linked mannosamine derivative (ManNDAz, **2**) (Scheme 1). ManNDAz (**2**) was then used as the substrate in a one-pot enzymatic reaction with recombinant *Escherichia coli* aldolase and *Neisseria meningitidis* CMP-Neu5Ac synthetase (CSS) to prepare the diazirine-modified CMP-Neu5DAz (**1**).^35^ While we found that the enzymes tolerated the unnatural diazirine to afford CMP-Neu5DAz with an overall yield of 54%, full conversion of ManNDAz (**2**) to the product **1** was not observed even after prolonged reaction times and addition of excess enzymes. The reaction was quenched after 5-6 h to minimize hydrolysis of **1** by the addition of alkaline phosphatase to degrade excess CTP, which also aided purification by size exclusion column chromatography over P2 Bio-Gel. In addition to preparing a CMP-Neu5DAz (**1**) substrate for enzymatic cell-surface remodeling, ManNDAz was also per-*O-*acetylated to provide the MOE probe Ac_4_ManNDAz to compare the efficiency of photo-crosslinking with our cell-surface glyco-engineering strategy.^15^

**Scheme 1.**
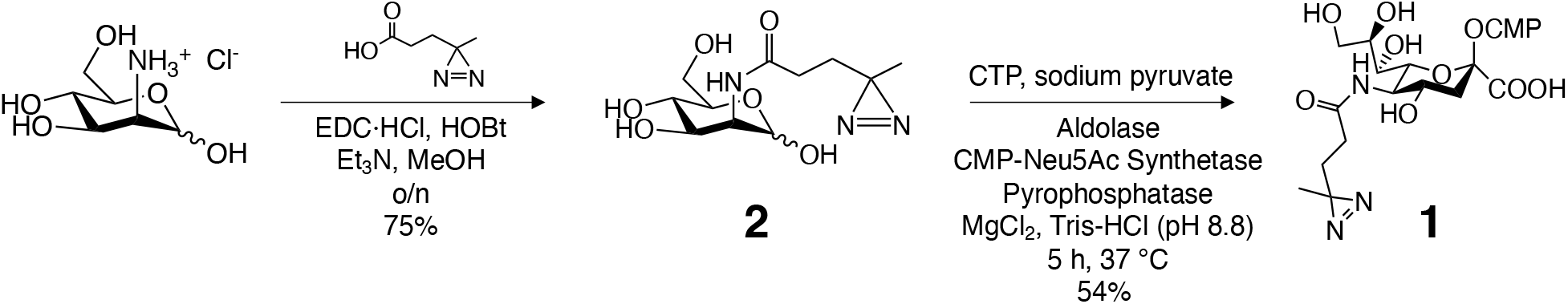
Chemo-enzymatic synthesis of diazirine-linked CMP-Neu5DAz (**1**).

### Cell-Surface Glycan Remodeling using CMP-Neu5DAz

With CMP-Neu5DAz (**1**) in hand, we employed exogenous enzymatic transfer to install this probe onto cell-surface glycoproteins. While one-step glyco-engineering protocols utilizing a concurrent neuraminidase treatment have improved labeling efficiencies in comparison to multi-step remodeling strategies, this is only feasible when bulky substituents are present at the C-5 position as these derivatives are not tolerated by many neuraminidases.^26-28^ The small diazirine-linker however, is insufficient to prevent hydrolysis by these enzymes.^13^ Thus, we employed a two-step approach where sialic acid termini on native cellular glycans were first cleaved by neuraminidase treatment, followed by remodeling with CMP-Neu5DAz and the sialyltransferase ST6GAL1.^16, 28^ We opted to use human ST6GAL1 as this enzyme could be transiently expressed in Expi293 cells with high yields.^40^ This enzyme has exceptionally high SEEL labeling efficiency, it selectively forms α-2,6-linked sialosides on *N*-glycans, and using the human sialyltransferase would recapitulate native α-2,6-sialylation which would more accurately represent native glycan-protein binding interactions.^27, 28, 37^

A panel of model cell lines were used in enzymatic glyco-engineering studies to demonstrate the broad applicability of our approach. This included adherent breast cancer cell lines SK-BR-3, MCF-7, and HS-578-T, and suspension U937 monocytes and RAJI B-cells. RAJI cells were selected as they overexpress CD22 (Siglec 2),^41^ which recognizes α-2,6-linked sialic acid on several glycoproteins, including other CD22 glycoproteins in *cis*-interactions.^13^ These interactions have been captured and identified by photo-crosslinking probes introduced on cell-surface glycans through MOE.^13-15^ We therefore reasoned that CD22 would serve as an ideal target to demonstrate proof-of-principle photo-crosslinking through an exo-enzymatic glycan remodeling strategy.

Cells were first desialylated by treatment with *Clostridium perfringens* sialidase (NanH) for 30 min. Next, α-2,6-linked Neu5DAz was installed by treating desialylated cells with 200 μM CMP-Neu5DAz and 20 μg/mL ST6GAL1 for 2 hours. Cell surface sialic acid was detected by flow cytometry using *Sambucus nigra* agglutinin (SNA), a lectin that recognizes α-2,6-linked sialosides, the epitope produced by ST6GAL1. Untreated cells generally displayed high binding to SNA, indicating high native levels of α-2,6-sialylation, and SNA binding was decreased following NanH treatment (Figure 2). SNA binding to HS-578-T and MCF-7 cell lines only decreased minimally after NanH treatment which may reflect low levels of native α-2,6-sialylation. Desialylation of these cell lines following NanH treatment (Figure 2b,c) was therefore confirmed by decreased *Maackia amurensis* lectin-II (MAL-II) binding, a lectin which prefers α-2,3-linked sialosides. (Figure S2). After remodeling desialylated cells with CMP-Neu5DAz and ST6GAL1, SNA binding was restored (SK-BR-3s, RAJIs; Figure 2a,d) or was higher (MCF-7s, HS-578-Ts, U937s; Figure 2b,c,e) than SNA binding to control cells, demonstrating efficient incorporation of the Neu5DAz probe on to cell-surface glycoconjugates. Glycan labeling of desialylated cells in the absence of either ST6GAL1 or CMP-Neu5DAz did not drastically restore SNA binding in any of the cell lines investigated (Figure S1). Small changes in SNA binding to RAJI cells observed in the absence of ST6GAL1 may be attributed to endogenous extracellular sialyltransferases, which have previously been identified in B-cells (Figure 3a, Figure S1).^42^

**Figure 2.**
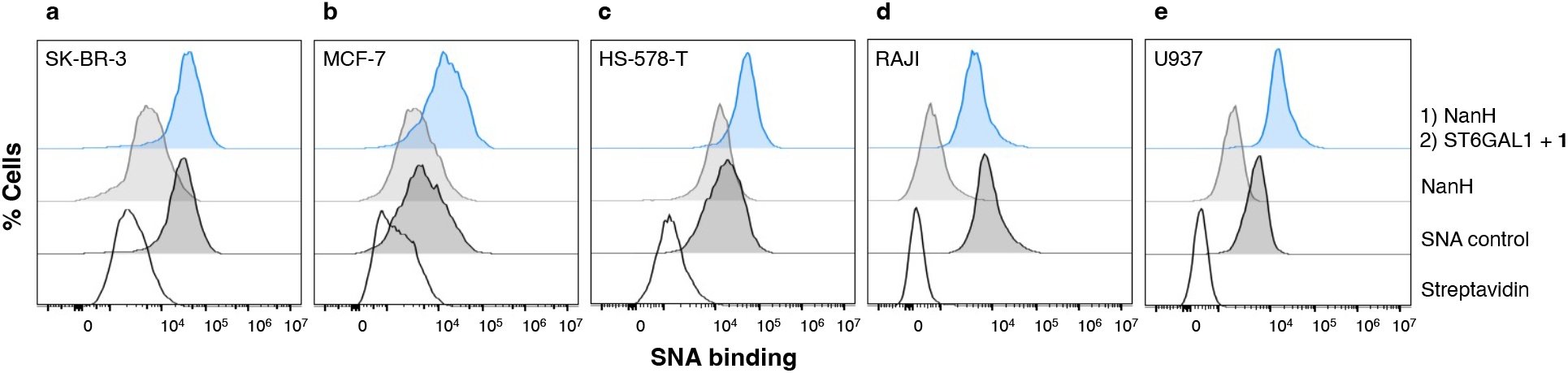
Cell-surface remodeling and display of CMP-Neu5DAz. Cells were labeled with CMP-Neu5Daz (**1**) using a two-step protocol. Cells were desialylated with sialidase (NanH) for 30 min at 37°C, followed by remodelling with 20 μg/mL ST6GAL1 and 200 μM **1** for 2 h at 37°C. Labeling with **1** on live cell surfaces was assessed by flow cytometry. Cells were stained with biotinylated SNA followed by Streptavidin-Pacific Blue, and were co-stained with PI to exclude non-viable cells.

**Figure 3.**
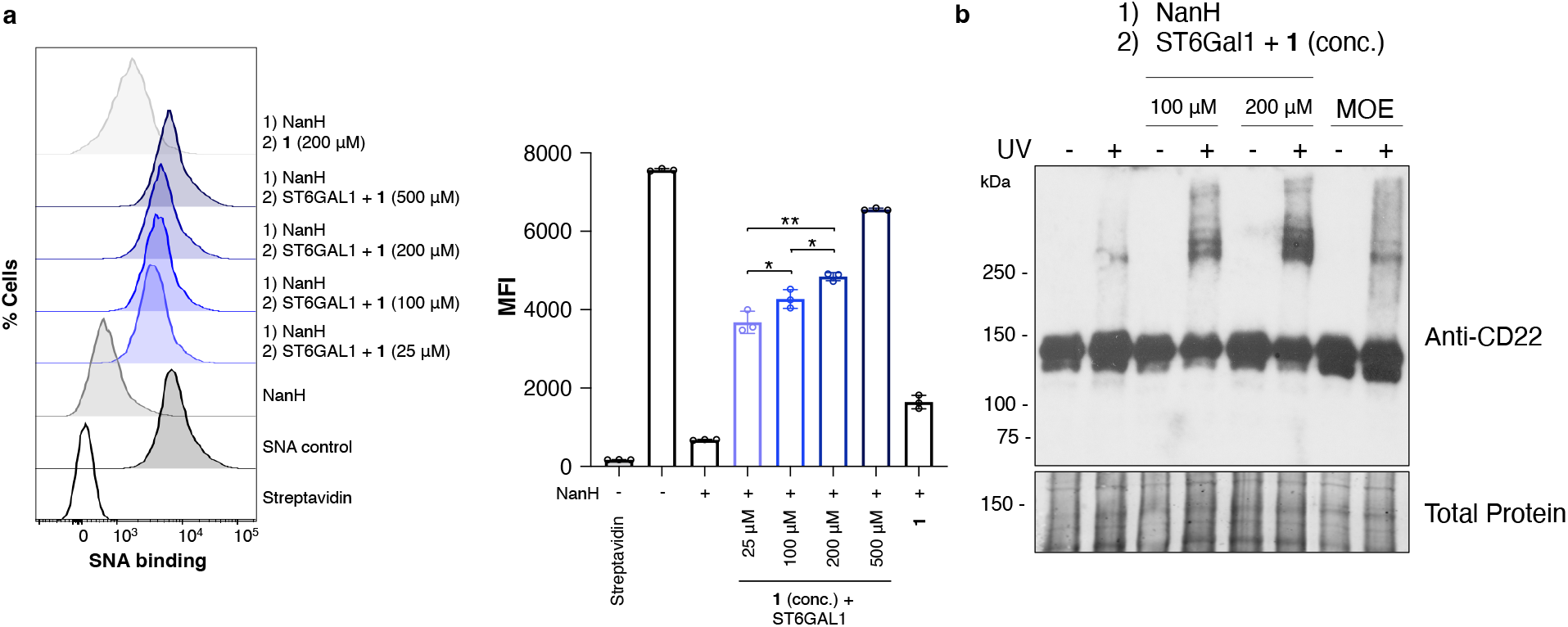
Cell-surface glycan labeling with CMP-Neu5DAz enables photo-crosslinking. **A)** Concentration-dependent labeling of live RAJI cells with ST6GAL1 and CMP-Neu5DAz (**1**) assessed by SNA binding by flow cytometry. Cells were stained with biotinylated SNA, detected with Streptavidin-Pacific Blue, and co-stained with PI to exclude non-viable cells prior to analysis. Median fluorescence intensity (MFI) of each sample was calculated. Error bars represent the standard deviation of three replicates (n = 3). Statistical significance was assessed by unpaired t-test with * p < 0.05 and ** p < 0.01. **B)** Photo-crosslinking of CD22-interaction complexes. RAJI cells were enzymatically remodeled by NanH followed by CMP-Neu5DAz (**1**) and ST6GAL1, or by MOE using 100 μM Ac4ManNDAz for 72 hrs. After labeling, cells were UV irradiated, lysed, and analyzed by immunoblot using an anti-CD22 antibody.

Next, we examined whether the amount of **1** displayed on the cell-surface RAJI cells could be modulated as this could impact binding and photo-crosslinking of CD22. Desialylated RAJI cells were remodeled with ST6GAL1 and various concentrations of CMP-Neu5DAz (**1**), and Neu5DAz incorporation was assessed by flow cytometry using SNA. As expected, the extent of cell-surface probe display was dependent on the concentration of CMP-Neu5DAz (**1**) used in the glyco-engineering step (Figure 3a).^26, 29^ Notably, the highest degree of labeling was obtained when 500 μM CMP-Neu5DAz was used and 25 μM CMP-Neu5DAz was sufficient for efficient incorporation on RAJI cells. These results indicate that the extent of probe display can be tuned. This is an attractive feature of the exo-enzymatic labeling strategy as the density of glycan epitopes can play an important role mediating binding to certain glycan-binding proteins,^8-11^ and this level of control cannot be easily obtained through MOE.^21, 22^

### Photo-crosslinking of CD22 *cis*-interaction complexes

Having demonstrated that exo-enzymatic glycan engineering could be applied to install Neu5DAz on multiple cell lines, including RAJI B-cells which overexpress CD22, we next sought to examine the photo-crosslinking capability of the installed probed. After desialylation of RAJI cells using NanH, cells were subsequently exo-enzymatically labeled with ST6GAL1 and 100, 200 or 500 μM CMP-Neu5DAz (**1**). Immediately after remodeling, cells were UV irradiated at 365 nm for 20 min to activate the diazirine probe and covalently capture α-2,6-sialoside-dependent glycan-protein interactions. Photo-crosslinked CD22-glycan complexes were detected by immunoblotting (IB) using an anti-CD22. Crosslinking was observed at all concentrations of CMP-Neu5DAz (**1**) assessed as indicated by the presence of higher molecular weight CD22-positive crosslinked bands (Figure 3, Figure S4). A small amount of background crosslinking was detected in control RAJI cells that were UV irradiated but not enzymatically remodeled. Crosslinking was dependent on UV activation and on the presence of both ST6GAL1 and CMP-Neu5DAz (**1**); in the absence of sialyltransferase or nucleotide-sugar probe, non-specific crosslinking was comparable to RAJI control cells (Figure S3). To compare crosslinking efficiency with MOE, RAJI cells were also incubated with 100 μM Ac_4_ManNDAz for 72 h and crosslinked.^15, 43^ In comparison to MOE, photo-crosslinking was enhanced when exo-enzymatic glycan labeling was used to install Neu5DAz as indicated by more intense higher molecular weight bands (Figure 3b).

Next, we examined the functional persistence of the exo-enzymatically installed Neu5DAz probe by assessing crosslinking efficiency after various incubation times following glycan remodeling. While all three CMP-Neu5DAz (**1**) concentrations assessed demonstrated crosslinking, we opted to use 200 μM CMP-Neu5DAz as this concentration had robust cell-surface labeling indicated by lectin staining (Figure 3a) and was most efficient for photo-crosslinking (Figure S4). After remodeling desialylated RAJI cells with ST6GAL1 and CMP-Neu5DAz (**1**), Neu5DAz-engineered cells were cultured in medium for 1, 3, or 6 h prior to UV irradiation (Figure 4). Crosslinking efficiency 1 h after enzymatic labeling was comparable to when Neu5DAz-engineered cells were UV irradiated immediately following remodelling (0 h incubation). Crosslinking could still be detected up to 6 hours following enzymatic Neu5DAz installation, although with reduced efficiency compared to shorter incubation times. The time dependence on crosslinking efficiency may be attributed to a more rapid turnover of some labeled glycoproteins after longer incubation times,^26, 27, 44^ or recycling of CD22 which could disrupt the Neu5DAz-glycan-CD22 complexes prior to crosslinking.^45^ However, as crosslinked complexes were still detectable up to 6 h after displaying Neu5DAz on cell-surfaces, this method may prove useful for studying interactions are slower to form.

**Figure 4.**
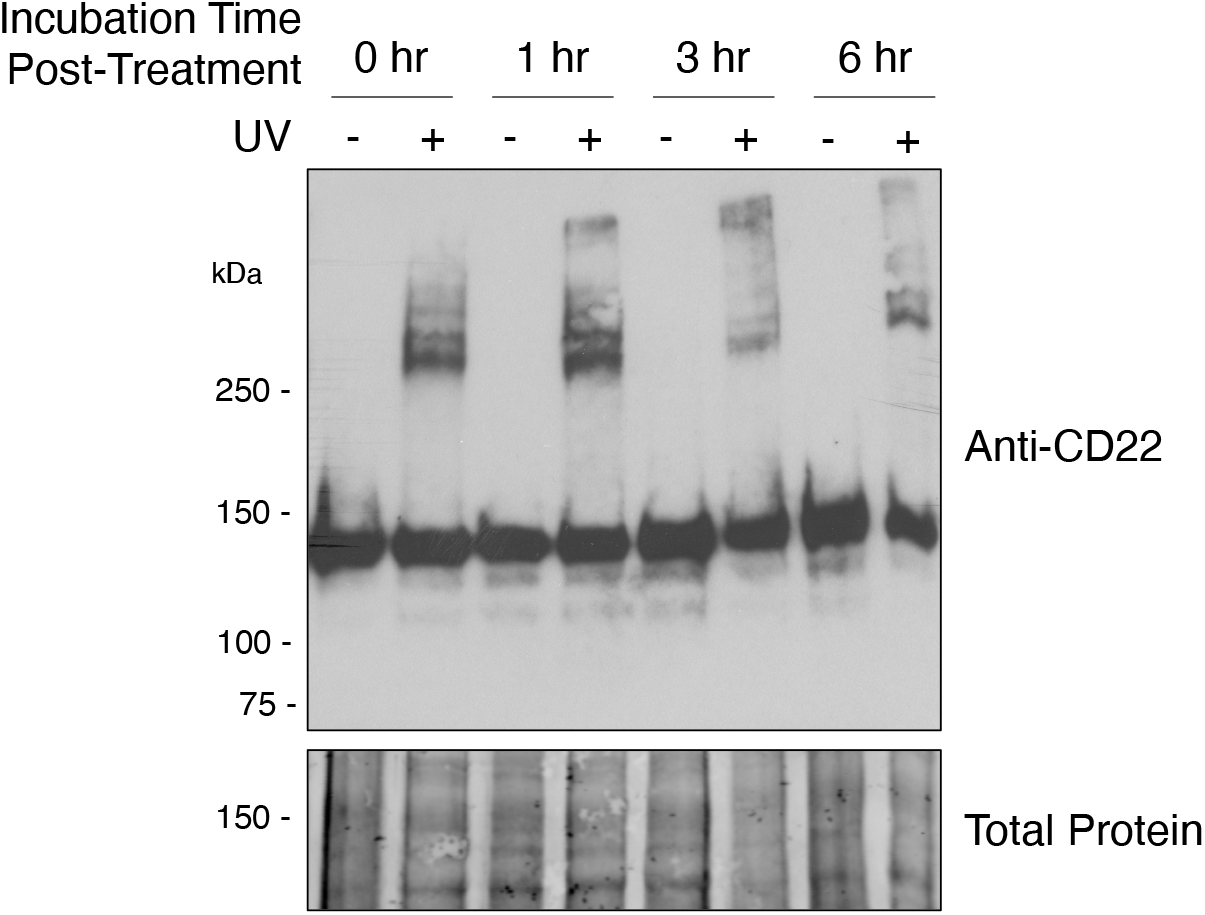
Functional persistence of exo-enzymatically installed Neu5DAz photo-crosslinker. After remodeling RAJI cells with ST6GAL1 and CMP-Neu5DAz (**1**), cells were incubated in culture medium for the indicated time prior to UV irradiation and analysis by immunoblot using an anti-CD22 antibody.

## Conclusions

Photo-crosslinking diazirine-linked probes have previously been incorporated into cellular glycans through metabolic oligosaccharide engineering (MOE).^15-18, 20^ Here, we demonstrate that exo-enzymatic cell-surface glycan labeling can be used as an efficient method to introduce a photo-crosslinking sialic acid on to cellular glycoproteins. A two-step approach was employed where native cellular sialic acid is first cleaved by a neuraminidase NanH, followed by selective α-2,6-Neu5DAz introduction onto *N*-glycans using CMP-Neu5DAz and the sialyltransferase ST6GAL1. Neu5DAz probe installation was demonstrated on multiple cell lines, density could be controlled by altering the concentration of CMP-Neu5DAz, and photo-crosslinked *cis*-CD22 interaction complexes on RAJI cells were observed over various probe concentrations and times post-glycan remodeling. Our method is highly complementary to an approach recently reported by the Kohler group that was simultaneously preprinted while finalizing our studies reported here. In their report, they use an *A. ureafaciens* sialidase and a bacterial α-2,6-sialyltransferase from *P. damsela* to engineer different cell lines than those used in this report with a CMP-Neu5DAz derivative, highlighting the versatility of this exo-enzymatic.^46^

An exo-enzymatic approach is advantageous over an MOE strategy as the photo-crosslinking sugar can be installed on cells in a few hours, it is broadly applicable to various cell types, and crosslinking was enhanced compared to MOE. Furthermore, this method offers more control over selective installation of the crosslinking sugar as well-defined glycan linkages, epitopes, and selective glycan subclass labeling can be achieved by harnessing the substrate specificity of the glycosyltransferase used. This work expands the scope of methodologies available to incorporate diazirine-based photo-crosslinking sugars on to cellular glycans and lays the foundation for exploring directed exo-enzymatic glycan labeling to interrogate and characterize glycan-protein interactions. We envision that the CMP-Neu5DAz probe could be used with other sialyltransferases enzymes and that analogous strategies could be developed to exo-enzymatically label glycans with other photo-crosslinking nucleotide-sugar probes (eg. UDP-GlcNAc, UDP-GalNAc). Selective display of photo-crosslinking probes on cell-surface glycans broadens the tools available to study and characterize glycan-protein interactions and will facilitate the identification of binding partners involved in these critical interactions.

### Experimental Section

#### Materials

Biotinylated *Sambucus nigra* agglutinin (SNA) (cat # B1305) and *Maackia amurensis* lectin-II (MAL-II) (cat # B1265) were purchased from Vector labs. Streptavidin-PacificBlue conjugate (cat # S11222), rabbit monoclonal anti-CD22 antibody (cat # EP498Y) were purchased from Abcam. Goat anti-rabbit antibody conjugated to horseradish-peroxidase (HRP) (cat # 65-6120) was purchased from Thermo-Fisher Scientific.

#### Cell Culture

RAJI and U937 cells were cultured in RPMI-1640 medium (with L-glutamine, sodium bicarbonate) supplemented with 10% fetal bovine serum (FBS) and 1X penicillin/streptomycin (P/S). Cells were passaged after reaching 2 × 106 viable cells/mL. SK-BR-3 and HS-578-T cells were cultured in high glucose DMEM medium with L-glutamine, supplemented with 10% FBS and 1X P/S. MCF-7 cells were cultured in MEM with Earle’s salts, L-glutamine, and sodium bicarbonate supplemented with 10% FBS and 1X P/S. Adherent cells were passaged using 0.25% trypsin-EDTA after reaching ∼80% confluency. All cells were maintained in a humid 5% CO_2_ atmosphere at 37 °C.

#### Cell-Surface Glycan Labeling

Adherent cells were plated in 12-well plates (300 000 cells/well) and grown to 80% confluency. Suspended cells were centrifuged and resuspended at 6 M cells/mL. Prior to labeling, cells were washed three times with culture medium without FBS. **Neuraminidase treatment:** Washed cells were incubated with 50 μg/mL *Clostridium perfringens* neuraminidase (NanH) in serum-free culture medium for 30 min and then were subsequently washed three times with serum-free culture medium. **Enzymatic Glycan Labeling:** Washed desialylated cells were incubated with a mixture of serum-free culture medium containing CMP-Neu5DAz (**1**), 0.1% BSA, and 20 μg/mL ST6GAL1 for 2 h at 37 °C. Untreated control experiments were treated with a mixture of serum-free culture medium containing 0.1% BSA with or without 20 μg/mL ST6GAL1 or 200 μM CMP-Neu5DAz. Following glycan labeling, cells were washed three times with 1% FBS/DPBS then treated as indicated.

#### Flow Cytometry Analysis

For the detection of cell-surface α-2,6-linked sialic acid and confirmation of sialic acid cleavage by NanH, control and enzymatically labeled adherent cells were first washed three times with DPBS without Ca/Mg and detached using 10 mM EDTA for 5 min at 37 °C. Cells were then suspended in 1% FBS/DPBS, centrifuged gently (300 rpm for 3 min), washed twice with 1% FBS/DPBS, and resuspended in 200 μL of staining buffer (1% FBS/DPBS) with 20 μg/mL biotinylated-SNA or biotinylated-MAL-II for 30 min at 4 °C. Cells were then washed/centrifuged once with staining buffer, then incubated with streptavidin-Pacific Blue (2.5 μg/mL in staining buffer) for 30 min at 4 °C in the dark. The cells were washed once in staining buffer, centrifuged gently and resuspended in 300 μL of FACS buffer (PBS without Ca/Mg supplemented with 2 mM EDTA and 0.5% BSA) and transferred to 96-well plates for flow cytometric analysis (Beckman Coulter, Cytoflex S). Cell viability was determined by adding 1 μg/mL PI to cell suspensions 1 min prior to analysis. The live population of cells was gated based on forward and side scatter emission, and exclusion of PI positive cells on the FL11 (610/20 BP filter) emission channel. Streptavidin-Pacific Blue binding was determined by fluorescence intensity on the FL6(450/45 BP filter) emission channel.

#### Photo-Crosslinking

After cell-surface glycan labeling, cells were either washed and allowed to incubate for various timepoints (1, 3, or 6 h) in complete medium at 37 °C, or immediately washed three times in DPBS. Cells were resuspended in 1 mL DPBS in a 12-well plate. Plates were set on ice and placed 1 cm away from the light source and irradiated at 365 nm with a UV lamp (Prolume F8T5BLB) for 20 min. Cells were then transferred to Eppendorf tubes and gently washed/centrifuged once in cold DPBS.

#### Immunoblot Blot Analysis

After photo-crosslinking, cells were lysed with RIPA lysis buffer containing 1X protease inhibitor cocktail (NEB cat # 5871S). Cell lysates were clarified by centrifugation at 14000g for 15 min and the total protein content of the supernatants was assessed by BCA assay (Pierce, ThermoFisher). Samples (30 μg total protein) were were resolved on a 6% SDS-PAGE gel and transferred to a low-fluorescence PVDF membrane (Immobilon-FL, Sigma). For normalization, total protein was stained with Revert Total Protein stain (Li-Cor Biosciences) for 5 min, washed twice with wash buffer (6.7% acetic acid, 30% methanol), and rinsed briefly in TBS prior to scanning on an Odyssey Li-Cor CLx scanner (Li-Cor Biosciences). Next, the membrane was blocked in blocking buffer (5% nonfat dry milk in TBS) for 1 h at room temperature. The blocked membrane was incubated for 1.5 h at room temperature with anti-CD22 antibody (1:5,000) in blocking buffer with 0.1% Tween-20 (TBST) and washed with TBST (6 × 10 min). The membrane was then incubated for 1 h at room temperature with goat anti-rabbit antibody conjugated to HRP (1:25,000) in TBST, followed by washing (3 × 10 min) with TBST. Membranes were rinsed briefly in TBS prior to imaging. Final detection of HRP activity was performed using SuperSignal West Atto Ultimate Sensitivity Substrate (ThermoFisher), exposure to film (Amersham) and development using a digital X-ray imaging machine.

## Acknowledgements

The authors thank Dr. Matthew Macauley (University of Alberta) for providing HS-578-T and U937 cell lines. We acknowledge the New Frontiers in Research Fund (SSHRC-NFRF), the National Sciences and Engineering Research Council of Canada (NSERC) and the Canada Foundation for Innovation (CFI) for funding supporting this work. J.L.B. acknowledges NSERC for a CGS-M scholarship and Queen’s University for an E. G. Bauman Fellowship award.

