## Supplementary Information for "Exo-Enzymatic Cell-Surface Glycan Labeling for Capturing Glycan-Protein Interactions through Photo-Crosslinking"

#### 1. Supporting Figures

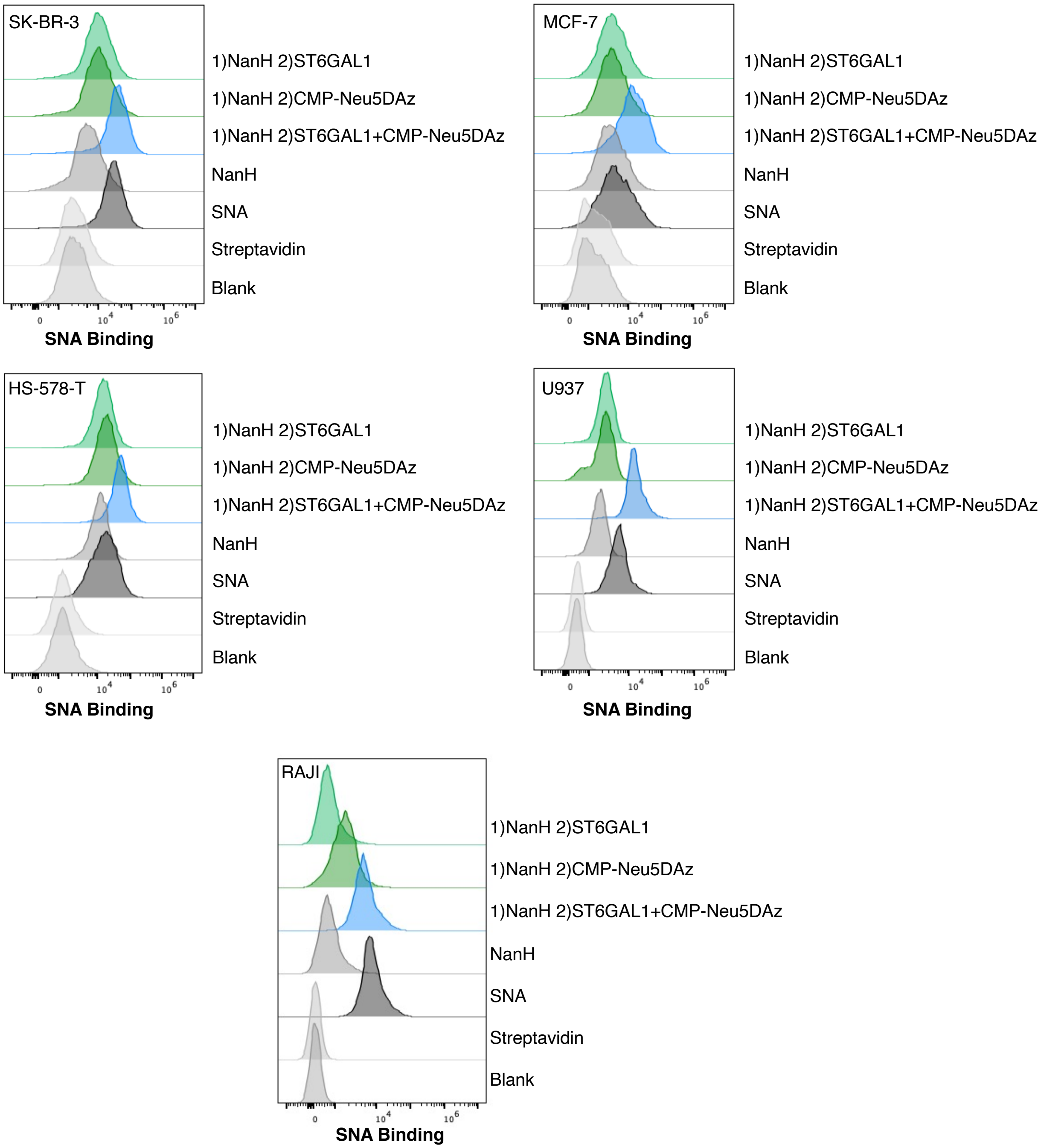


**Figure S1. Enzymatic cell-surface glycan labeling with CMP-Neu5DAz (1) and ST6GAL1.** Cells were labeled with CMP-Neu5Daz (**1**) using a two-step protocol. Cells were desialylated with sialidase (NanH) for 30 min at 37°C, followed by remodeling with or without 20 µg/mL ST6GAL1 and 100 µM **1** for 2 h at 37°C. RAJI cells were labeled with 200 µM CMP-Neu5DAz. Cells were stained with biotinylated SNA followed by Streptavidin-Pacific Blue, and were co-stained with PI to exclude non-viable cells.

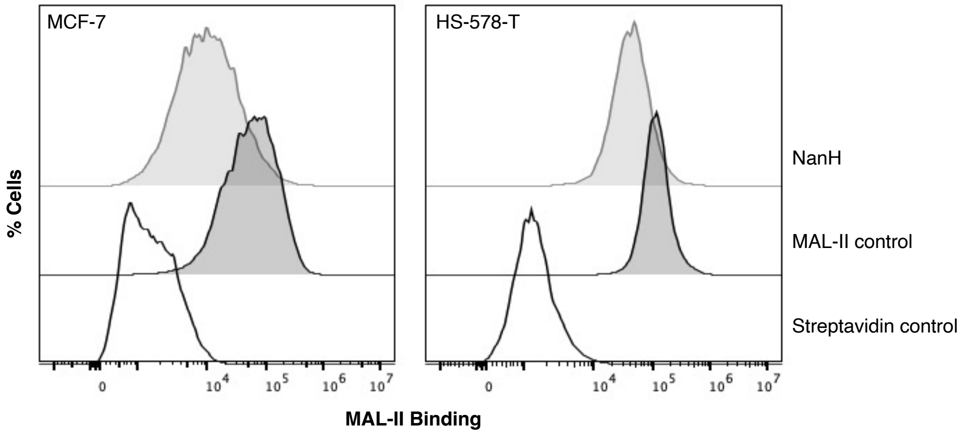


**Figure S2. Monitoring sialidase cleavage with MAL-II.** MCF-7 and HS-578-T cells were desialylated with sialidase (NanH) for 30 min at 37 °C. Sialic acid cleavage was assessed by flow cytometry. Cells were stained with 20 µg/mL biotinylated MAL-II followed by Streptavidin-Pacific Blue, and were co-stained with PI to exclude non-viable cells.

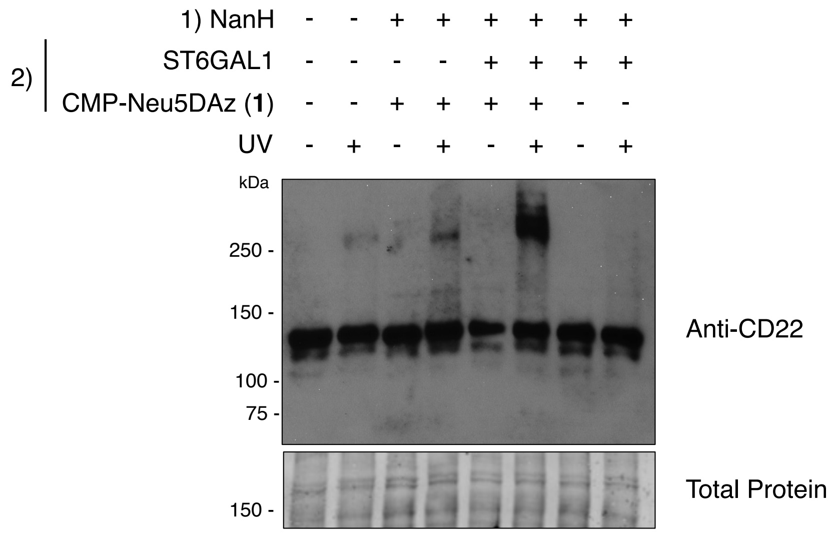


**Figure S3. Photo-crosslinking is dependent on the addition of ST6GAL1 and CMP-Neu5DAz (1).** RAJI cells were enzymatically desialylated NanH and subsequently labeled with or without 200 µM CMP-Neu5DAz (**1**) and ST6GAL1. After labeling, cells were UV irradiated, lysed and analyzed by immunoblot using an anti-CD22 antibody.

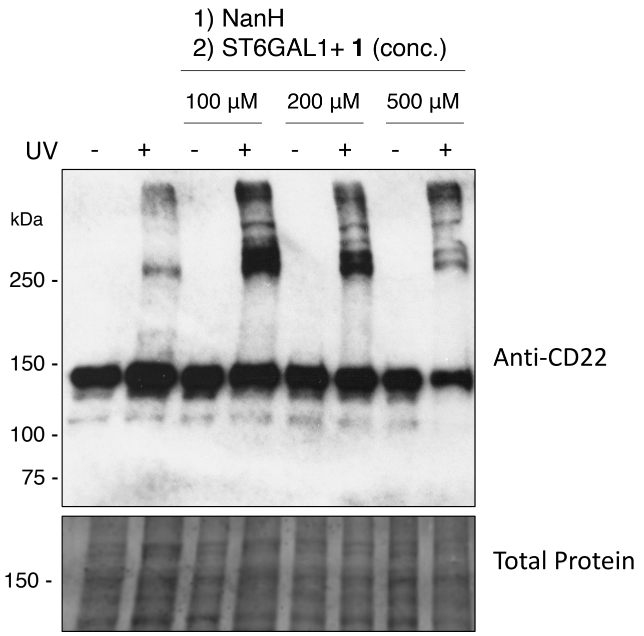


**Figure S4. Concentration dependence of CMP-Neu5DAz for photo-crosslinking.** RAJI cells were enzymatically remodeled by NanH followed by various concentrations of CMP-Neu5DAz (**1**) and ST6GAL1. After labeling, cells were UV irradiated, lysed, and analyzed by immunoblot using an anti-CD22 antibody.

#### 2. Chemo-Enzymatic Synthesis

##### Cytidine-5'-monophospho-5-*N*-[4,4-azo-pentamido]-3, 5- dideoxy-β-D-glycero-D-galacto-2-nonulopyranosonic acid (CMP-Neu5DAz, 1):

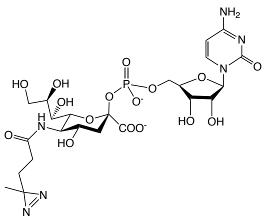


ManNDAz (**2**) was prepared as previously described in the literature.[1-3] ManNDAz (50 mg, 0.17 mmol), sodium pyruvate (95.06 mg, 0.86 mmol), and CTP disodium salt (118.4 mg, 0.22 mmol) were dissolved in a Tris–HCl buffer (0.1 M, 17.2 mL, pH 8.8) containing $MgCl_2$ (20 mM). To this mixture, recombinant *E. coli* aldolase (35 mg/mmol substrate), *N. meningitidis* CMP-sialic acid synthetase (31 mg/mmol substrate) and *P. multocida* inorganic pyrophosphatase (18 mg/mmol substrate) were added and the reaction was incubated at 37 °C with shaking (200 rpm), in the dark. Reaction progress was monitored by TLC (7:3 EtOH/1 M aqueous ammonium bicarbonate), which was allowed to proceed until disappearance of the ManNDAz starting material (Rf = 0.8), detected by staining with *p*-anisaldehyde. The reaction was then quenched by the addition of calf intestinal alkaline phosphatase (CIAP; NEB cat # M0525) to degrade unreacted CTP to aid subsequent purification steps. The mixture with CIAP was incubated at 37 °C with shaking until CTP was no longer visible by TLC (detected by UV exposure; Rf = 0.1), which otherwise co-elutes with the product over P2-BioGel purification. Precipitates were removed by centrifugation and enzymes were then removed by spin-filtration (10 kDa MWCO, Amicon Ultra-15 Centrifugal Filter, cat # UFC9010) before flash-freezing with liquid $N_2$ and lyophilization. The isolated residue was resolubilized in a 50 mM ammonium bicarbonate buffer and loaded on a fine P2-BioGel® (Bio-Rad) size exclusion column. Fractions containing the title compound were collected on ice in the dark, pooled, and lyophilized to afford CMP-Neu5DAz (**1**) as a white powder (64 mg, 54%). CMP-Neu5DAz (**1**) was stored as a dry solid at -80 °C and was dissolved in culture medium immediately before use. $^1$H NMR (700 MHz, $D_2O$) δ 7.99 (d, *J* = 7.5 Hz, 1H, H-6 cyt), 6.14 (d, *J* = 7.5 Hz, 1H, H-5 cyt), 6.00 (d, *J* = 4.6 Hz, 1H, H-1 rib), 4.35 – 4.27 (m, 2H, H-2 + H-3 rib), 4.28 – 4.20 (m, 3H, H-5a, H-5b, H-4 rib), 4.17 (d, J = 10.4 Hz, 1H, H-6), 4.08 (td, *J* = 10.7, 4.5 Hz, 1H, H-4), 4.02 – 3.88 (m, 3H, H-5, H-8, H-9a), 3.61 (dd, *J* = 11.8, 6.7 Hz, 1H, H-9b), 3.54 (d, *J* = 9.4 Hz, 1H, H-7), 2.50 (dd, *J* = 13.2, 4.6 Hz, 1H, $H\text{-}3_{eq}$), 2.23 (td, *J* = 11.9, 5.8 Hz, 2H, $C\underline{H}_2$CO-DAz), 1.85 – 1.60 (m, 3H, $H\text{-}3_{ax}$ + $C\underline{H}_2CN_2$-DAz), 1.04 (s, 3H, $C\underline{H}_3CN_2$-DAz). $^{13}$C NMR (175 MHz, $D_2O$) δ 141.7, 96.6, 88.9, 83.1, 74.4, 71.8, 69.8, 69.2, 68.9, 66.9, 64.7, 63.1, 51.2, 41.2, 30.2, 29.9, 18.6. ESI-MS m/z calcd for $C_{23}H_{34}N_6O_{16}P$, $[M-H]^-$: 681.1774, found, 681.1798.

#### 3. Biological Procedures

##### Metabolic Oligosaccharide Engineering (MOE) using $Ac_4$ManNDAz

$Ac_4$ManNDAz was prepared and used in MOE experiments as previously described.[1, 2] RAJI cells were plated in a 10-cm dish (1 M cells/mL) and were grown in complete culture medium supplemented with 100 µM $Ac_4$ManNDAz for 72 h. Cells were then washed three times with DPBS and crosslinked as described.

##### Bacterial Protein Expression

Bacterial proteins *P. multocida* strain Pm70 inorganic pyrophosphatase (PmPpA), *E. coli* K12 sialic acid aldolase (aldolase), and *N. meningitidis* CMP-sialic acid synthetase (CSS) were recombinantly expressed as previously reported.[4, 5] The genes encoding aldolase and PmPpA were commercially synthesized and inserted into pET-15b vectors using NcoI and BamHI or NdeI and BamHI restriction sites, respectively (Genscript). The gene encoding *N. meningitidis* CMP-sialic acid synthetase (CSS) was commercially synthesized and inserted into a pET-22b(+) vector using NdeI and XhoI restriction sites (Genscript). The gene sequence of *C. perfringens* a2-3,-6,-8 neuraminidase (NanH) (GenBank: Y00963.1) was commercially synthesized and ligated into a pET-15b plasmid using NdeI and XhoI restriction sites (Genscript). The vectors containing the genes were chemically transformed into BL-21 competent cells. Transformed cells were cultured in 5 mL LB media supplemented with 100 µg/mL ampicillin for 16 h at 37 °C with shaking (200 rpm) for 16 hours. Cultured cells were then scaled up to 500 mL in LB-amp to an optical density ($OD_{600}$) of 0.8. Protein expression was then induced with addition of isopropyl β-D-thiogalactopyranoside (IPTG) to a final concentration of 0.1 mM and cells were incubated at 25 °C with shaking (200 rpm) for 16 hours before harvesting by centrifugation. Pelleted cells were resuspended in lysis buffer (0.1 M Tris-HCl, pH 8.0, 0.1% Triton X-100) and lysed by high pressure homogenization using an Avestin Emulsiflex C3. Cell debris was removed by ultracentrifugation, and the supernatant was mixed 1:1 with binding buffer (10mM imidazole,

500mM NaCl, 50mM Tris-HCl, pH 7.5). The resulting solutions were then filtered using a 0.22 µm Filter Unit and applied to a $Ni^{2+}$-sepharose resin (GE Healthcare, fast flow) pre-equilibrated with binding buffer. The column was washed with 6 column volumes (CV) of binding buffer, 6 CV wash buffer (50mM imidazole, 500mM NaCl, 50mM Tris–HCl, pH 7.5), and eluted with 10 CV elution buffer (200mM imidazole, 500mM NaCl, 50mM Tris-HCl, pH 7.5). Overexpression and purification were assessed by SDS-PAGE. Purified protein was buffer exchanged into 20 mM Tris-HCl (pH 7.5) buffer and concentrated using a 10 kDa MWCO spin filter (Amicon). All proteins were stored in 20 mM Tris-HCl (pH 7.5) with 20% glycerol at -80 °C, except for CSS which was stored in 20 mM Tris-HCl (pH 8.5) with 20% glycerol at -80 °C. Final protein concentrations were determined by BCA assay. Yields per 500 mL culture were: NanH = 175 mg; aldolase = 40 mg; CSS = 45 mg; PmPpA = 55 mg.

**Mammalian Protein Expression**

The plasmid encoding a soluble, secreted GFP fusion protein containing the catalytic domain of human β-galactoside α-2,6-sialyltransferase 1 (ST6GAL1) in a pGEn2-DEST vector was purchased from DNASU (cat # HsCD00413052). Recombinant GFP-ST6GAL1 protein was expressed and purified as previously described using Expi293 cells.[6] Plasmids were chemically transformed into DH5-alpha competent cells. Transformed cells were cultured in 5 mL LB media supplemented with 100 µg/mL ampicillin for 16 h at 37 °C and shaking (200 rpm) for 16 hours. Cultured cells were then scaled up to 500 mL and plasmids were isolated using the PureLink™ HiPure Plasmid Maxiprep Kit (Thermofisher, cat # K210006).

**Expi293 cell maintenance:** Expi293 cells (ThermoFisher) were cultured in Expi293 Expression Medium (ThermoFisher). Cells were maintained in a humid 5% $CO_2$ atmosphere at 37 °C with shaking at 120 rpm. Cells were passaged for maintenance after reaching $4 \times 10^6$ viable cells/mL. Cells were cultured for at least 3 passages following thaw prior to transfection.

**Transfection and Expression:** Expi293 cells were transiently transfected with the vector containing the ST6GAL1 gene using the Expifectamine 293 Transfection Kit (ThermoFisher, cat # A14524). On day 5 following transfection, cells were harvested by centrifugation (25 min, 4000 rcf). The supernatant was collected and adjusted to contain 20 mM imidazole, 200 mM NaCl, and 30 mM sodium phosphate, at pH 7.2. The resulting solutions were then filtered using a 0.45 µm PES Filter Unit and applied to a $Ni^{2+}$-sepharose resin pre-equilibrated with column buffer (20 mM HEPES, 300 mM NaCl, pH 7.2) containing 20 mM imidazole. The column was sequentially washed with column buffer containing 20 mM, 50 mM, and 100 mM imidazole prior to elution with 300 mM imidazole. Overexpression and purification were assessed by SDS-PAGE. Purified protein was buffer exchanged into 20 mM Tris-HCl (pH 7.5) buffer, concentrated using a 10 kDa MWCO spin filter (Amicon), and stored in 20 mM Tris-HCl (pH 7.5) with 20% glycerol at -80 °C. The final protein concentration was determined by BCA assay. Yield per 100 mL culture was 4.5 mg.

### 4. NMR Data

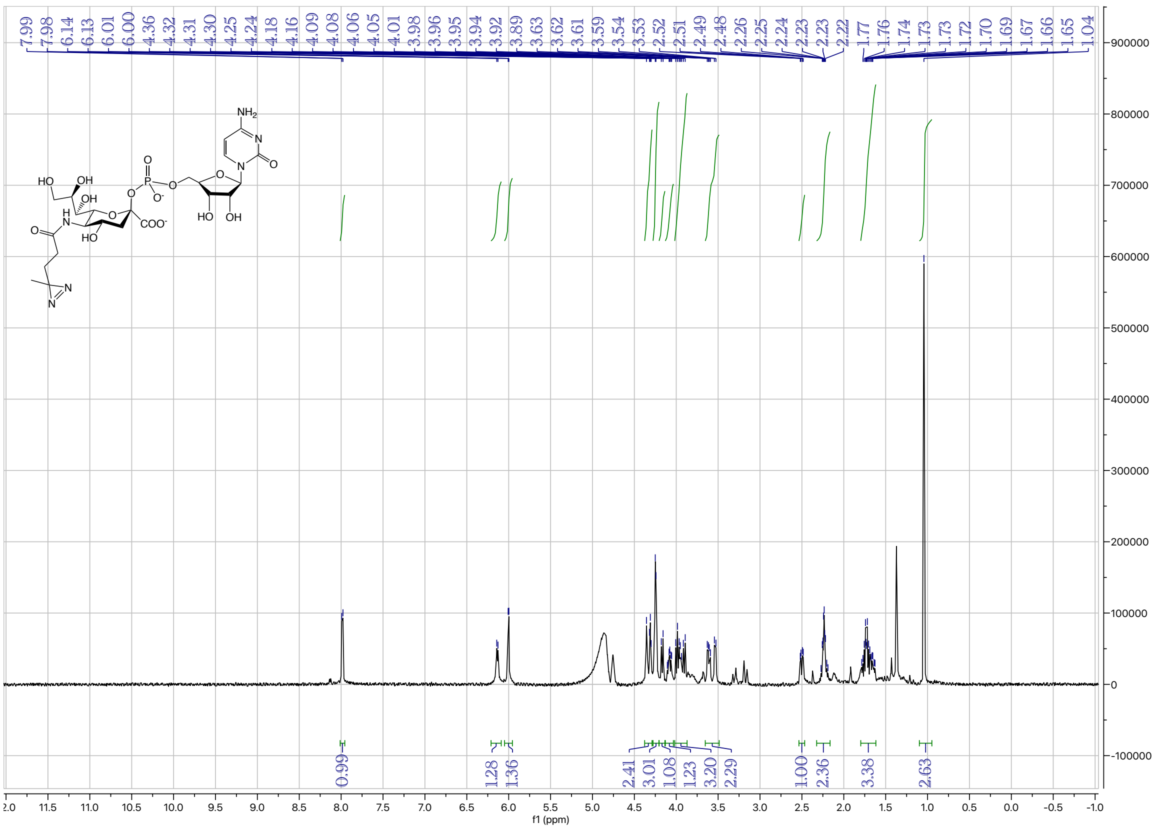

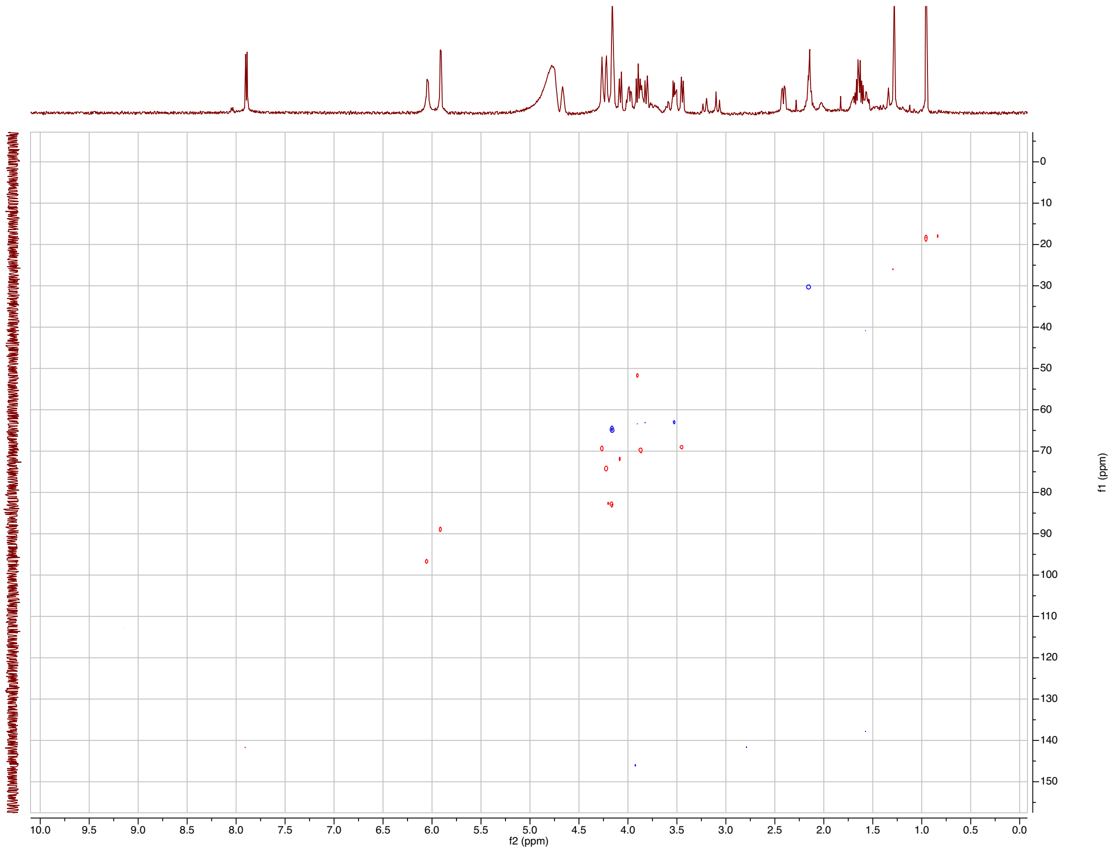
